# Random forest machine-learning algorithm classifies white- and brown-rot fungi according to the number of Carbohydrate-Active enZyme genes

**DOI:** 10.1101/2024.03.15.585254

**Authors:** Natsuki Hasegawa, Masashi Sugiyama, Kiyohiko Igarashi

**Affiliations:** Department of Biomaterial Sciences, Graduate School of Agricultural and Life Sciences, The University of Tokyo; Center for Advanced Intelligence Project, RIKEN; UT7 Next Life Research Group, The University of Tokyo

**Keywords:** Lytic polysaccharide monooxygenase, wood-rotting fungi, Carbohydrate-Active enZymes, machine learning, random forest algorithm

## Abstract

Wood-rotting fungi play an important role in the global carbon cycle because they are only known organisms that digest wood, the largest carbon stock in nature. In the present study, we used linear discriminant analysis and random forest (RF) machine learning algorithms to predict white- or brown-rot decay modes from the numbers of genes encoding Carbohydrate-Active enZymes (CAZymes) with over 98% accuracy. Unlike other algorithms, RF identified specific genes involved in cellulose and lignin degradation, including auxiliary activities (AA) family 9 lytic polysaccharide monooxygenases, glycoside hydrolase family 7 cellobiohydrolases, and AA family 2 peroxidases, as critical factors. This study sheds light on the complex interplay between genetic information and decay modes and underscores the potential of RF for comparative genomics studies of wood-rotting fungi.

**Importance:** Wood-rotting fungi are categorized as either white- or brown-rot modes based on the coloration of decomposed wood. The process of classification can be influenced by human biases. The random forest machine learning algorithm effectively distinguishes between white- and brown-rot fungi based on the presence of Carbohydrate-Active enZyme genes. These findings not only aid in the classification of wood-rotting fungi but also facilitate the identification of the enzymes responsible for degrading woody biomass.

## Introduction

The most abundant source of carbon on land is plant biomass, especially in forest ecosystems, where much of it is fixed in the form of dead wood(1). Wood cell walls are composed of three main components: cellulose, hemicellulose, and lignin. Cellulose is a linear homopolysaccharide composed of β-1,4-linked D-glucopyranose, and forms a rigid crystal structure through hydrogen bonding and hydrophobic interactions among the molecular chains(2). Hemicellulose is a heteropolysaccharide containing various sugar residues, including D-xylopyranose and D-mannopyranose, and generally consists of β-1,4-linked linear chains with short branching side chains. Lignin consists of complex and heterogeneous polymers formed by radical cross-linking of phenolic precursors, and is resistant to degradation(3). Cellulose makes up 40-50% of the dry weight of wood, while hemicellulose and lignin each make up 15-30%.

Wood-rotting fungi are unique in their ability to efficiently produce robust lignocellulosic enzymes that efficiently degrade these three components, and play an important role in the carbon cycle of forest ecosystems. Most wood-rotting fungi belong to the Basidiomycota, especially the Agaricomycetes and Dacrymycetes, although some, such as the soft-rot fungi, belong to the Ascomycota. These wood-rotting basidiomycetes are classified according to the two main decay patterns: white-rot fungi, which degrade all three components, and brown-rot fungi, which degrade only polysaccharides, cellulose and hemicelluloses (Fig. 1). However, a new type, called gray-rot fungi, has recently been reported to be intermediate between the two (4, 5). The degree of polymerization of cellulose gradually decreases during biomass decomposition by white-rot fungi, whereas it decreases rapidly in the early stages of decomposition by brown-rot fungi. In the case of brown-rot fungi, lignin is denatured but not degraded, leaving a macromolecular residue (6). These differences in the mode of wood decay have been supported in recent years by comparative genomics studies.

**Figure 1.**
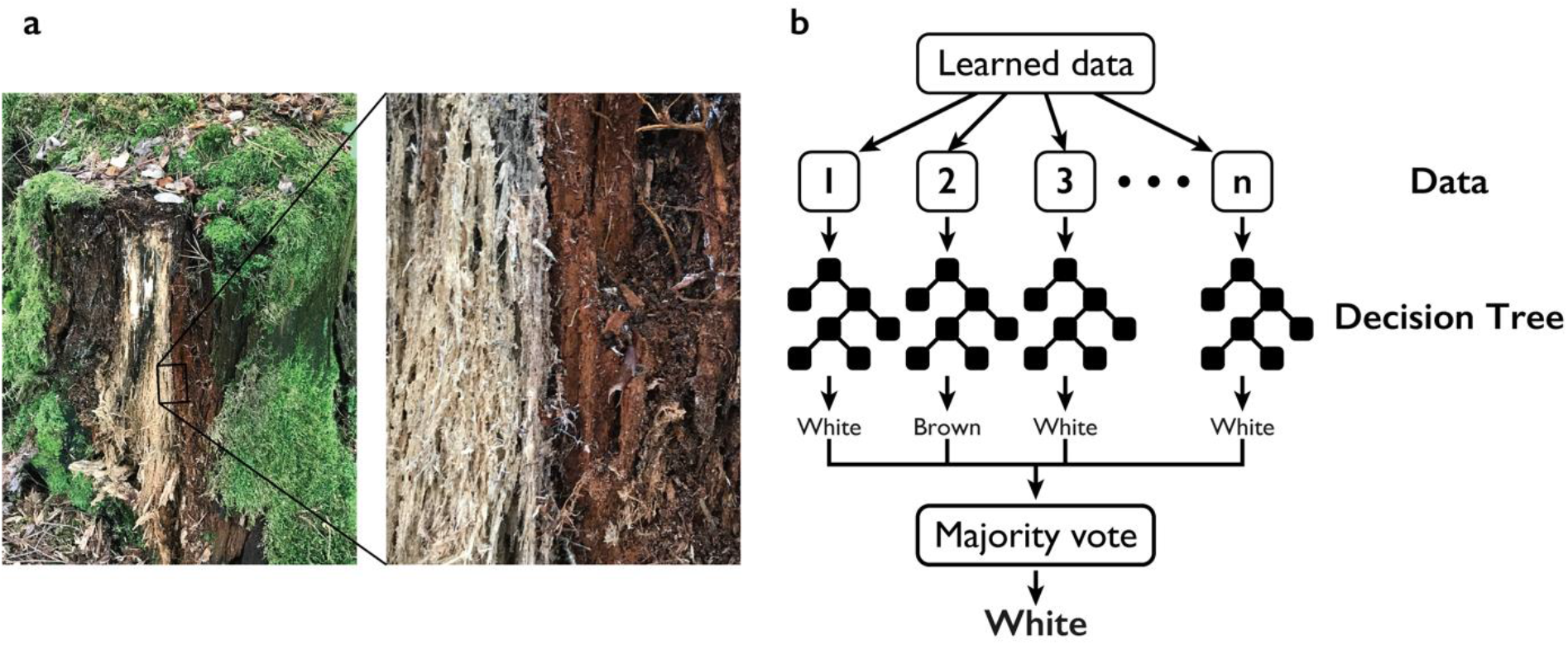
Typical white-rot and brown-rot decay modes seen in the forest and scheme of the random forest (RF) algorithm. (a) Two types of rotting (white- and brown-rot) are often observed even in the same log. (b) Scheme of the RF algorithm for the determination of wood-decay mode from the number of CAZymes.

The diverse Carbohydrate-Active enZymes (CAZymes) produced by wood-rotting fungi are classified into five classes in the CAZy database (http://www.cazy.org/) (7): glycoside hydrolases (GH), glycosyltransferases (GT), polysaccharide lyases (PL), carbohydrate esterases (CE), and enzymes with auxiliary activity (AA), in addition to the non-catalytic carbohydrate-binding modules (CBM). White-rot fungi generally possess more diverse genes than brown-rot fungi, especially in the enzyme families involved in the degradation of crystalline cellulose, such as GH6 and GH7, including cellobiohydrolase (CBH), and the AA9 family of lytic polysaccharide monooxygenases (LPMO) (8). AA2s, the class II peroxidase (POD) genes involved in lignin degradation, are also abundant in white-rot fungi and are present in low numbers or absent in the genome of brown-rot fungi(9). Thus, there is a consistent trend between the observed characteristics of each decay mode and the numbers of CAZymes genes in the fungus. An important advantage of comparative genomics, together with various omics analyses, is that we can infer how the fungus uses its enzymes, rather than simply examining what activity it has *in vitro*.

On the other hand, simple observational comparison of the numbers of genes is challenging. First, the number of enzyme families and genome samples that can be compared is limited, and if prior screening by the experimenter is required, this may lead to bias. Also, the observation of a limited number of samples makes it difficult to determine whether the differences found can be generalized as white-rot/brown-rot traits, and whether they are phylogenetic or random. Woody biomass has a complex structure with many different components, and many enzymes are involved in its decomposition in different ways. Further, in the CAZymes classification, it is not uncommon for enzymes with similar activities to be present in multiple families. Therefore, a more sophisticated mathematical method that can comprehensively evaluate the expression of diverse CAZymes genes in a wide range of wood-rotting fungal genomes is desirable.

One such multivariate analysis method is Fisher’s linear discriminant analysis (LDA)(10). More recently, machine learning has emerged as one of the best tools for analyzing multivariate data. In particular, the random forest (RF) strategy(11) has been proposed for machine learning in comparative genomics(12, 13). In this study, therefore, we applied these methods in order to gain insight into the genetic backgrounds of the white-/brown-rot modes of decay.

## Materials and Methods

### Generation of wood decay mode dataset

Most wood-rotting basidiomycetes fall into two orders, Agaricomycetes and Dacryomycetes, and CAZymes annotation data were obtained from MycoCosm (https://mycocosm.jgi.doe.gov/mycocosm/home)(14), the fungal genome database of the U.S. Joint Genome Institute (JGI), for all fungi classified into these two orders. Data were exported from the XML format to xlsx files; fungi without data for any of the six classes of CAZymes (AA, CBM, CE, GH, GT, and PL) were excluded from the dataset because it was unclear whether they lacked these genes or whether the annotation was incomplete. The resulting CAZymes annotation data included 455 basidiomycetes. These were searched for genera in the U.S. National Center for Biotechnology Information (NCBI) taxonomy database (https://www.ncbi.nlm.nih.gov/taxonomy) and assigned to an order level by examining the taxonomy of each fungus: *Intextomyces contiguus* (JGI project code: Intcon1_1), *Xenasmatella vaga* (Xenvag1), and *Sclerogaster hysterangioides* (sclhys1_1) were not assigned to an order level.

*Calocera viscosa* (Calvi1) was considered a white-rot fungus in the JGI MycoCosm but has been claimed to be a brown-rot fungus in some literature (15). It was treated as a brown-rot fungus in this study. *Sistotremastrum suecicum* (Sissu1) is also described as a brown-rot fungus in the JGI MycoCosm, but for the same reason(16, 17), it was treated as a white-rot fungus in this study. *Guyanagaster necrorhizus* (Guyne1) and *Cantharellales* sp. (Tvlas1) could not be identified by either method. Subsequent experiments were performed only with those fungi whose decay mode was identified here as white- or brown-rot. The CAZymes annotation data for these fungi is designated as the decay mode dataset.

### Creation of a phylogenetic tree

In the Markov clustering(18) function of JGI MycoCosm, all protein sequences of fungi belonging to Agaricomycotina are divided into 583,901 clusters. From these clusters, we selected those in which all fungi possessed at least one sequence, followed by the selection of the nine clusters from these clusters with the lowest average number of sequences possessed by each fungal species (Table 1).

**Table 1.** Gene clusters used for phylogenetic tree estimation.

| Cluster ID | Domain |
| --- | --- |
| 4393 | Ribosomal protein S7p/S5e |
| 4376 | Metallo-beta-lactamase superfamily |
| 4309 | Ribosomal protein L3 |
| 4137 | 50S ribosome-binding GTPase |
| 4099 | Chorismate synthase |
| 4019 | Proteolipid membrane potential modulator |
| 4014 | Mitochondrial carrier protein |
| 3993 | Pep3/Vps18/deep orange family |
| 3956 | Transport protein Trs120 or TRAPPC9, TRAPP II complex subunit |

BLAST searches on JGI MycoCosm were used to obtain the sequences of all homologous proteins of these nine clusters possessed by the fungi in the decay style dataset, and each cluster was multiply aligned using the MAFFT online service(19). The algorithm used was FFT-NS-i according to the manual recommendations; if a fungus possessed multiple copies of proteins of the same cluster, consensus sequences were generated on SeaView(20) after alignment. The resulting 9 multiple alignments were submitted to trimAI(21) on Phylemon 2.0(22), trimmed for poor alignments using the gappyout method, and concatenated.

Phylogenetic tree estimation was performed using the W-IQ-TREE web server(23). Multiple alignments were partitioned using Edge-linked, and substitution models were automatically selected using the model selection function, including FreeRate heterogeneity. Phylogenetic tree estimation was iterated 1000 times with ultrafast bootstrap.

### Choice of explanatory variables

In LDA analysis, it is difficult to obtain valid results when highly correlated combinations of explanatory variables (number of genes in each CAZy family/subfamily) exist in a dataset. This is because the importance of an explanatory variable to the model is more likely to be expressed as the importance of another variable correlated with it, rather than as the importance of the variable itself. This state of low independence among explanatory variables is called multicolinearity. To avoid this multicolinearity problem, the CAZy families were narrowed down using the following procedure.

First, a correlation ratio was calculated between the number of genes in each family and the white-rot/brown-rot trait, and those with a value of less than 0.1 were eliminated.

Then, for all remaining pairs of families with correlation coefficients greater than 0.55, the family with the lower correlation ratio to the trait was eliminated. After this refinement, the final 18 remaining CAZy families/subfamilies were subjected to a standardization treatment and used as explanatory variables in the LDA analysis (Table 2).

**Table 2.** CAZy families/subfamilies employed as explanatory variables in LDA.

| Explanatory variable | Correlated families (r) |
| --- | --- |
| AA1_1 | AA3 (0.63), AA3_2 (0.61), CE8 (0.61) |
| AA2 | — |
| AA3_1 | CE8 (0.59) |
| AA4 | — |
| AA5 | AA5_1 (1.00) |
| AA9 | AA8 (0.57), CBM1 (0.61), CE1 (0.62), CE15 (0.55), GH7 (0.60), GH10 (0.67), GH131 (0.62) |
| CE12 | CE8 (0.63), GH16 (0.59), GH35 (0.60), GH93 (0.58) |
| GH5_9 | GH16 (0.58) |
| GH15 | — |
| GH17 | — |
| GH45 | — |
| GH74 | — |
| GH79 | — |
| GH135 | — |
| GH145 | — |
| PL4_3 | — |
| PL8_4 | PL8 (1.00) |
| EXPN | — |

In the case of RF analysis, all CAZy families/subfamilies were used as explanatory variables because, in contrast to LDA, multicollinearity is not an issue. In addition, we did not standardize the explanatory variables because, in principle, the specific values of the explanatory variables are meaningless in the RF approach.

### LDA/RF Model Building and Evaluation

The decay style dataset was imbalanced as regards the number of samples of white- and brown-rot fungi. Since such imbalanced data set is likely to train a model that is biased toward the majority, we compared the performance of the model after training with a balanced dataset prepared by randomly generating hypothetical brown-rot fungus samples through oversampling. For oversampling, we used SMOTE(24) from the Python library imbalanced-learn.

The original or oversampled dataset was divided into training and test data in a 7:3 ratio. The training data was used to build the LDA/RF model and its performance was evaluated in terms of the percentage of correct responses, fit rate, recall rate, and F1 score against the test data (Fig. 2). For each test sample, LDA and RF models were built using the scikit-learn Python library tools, LinearDiscriminantAnalysis, or RandomForestClassifier. In addition, the coefficients of the discriminant function for LDA and the Gini importance for RF were obtained as measures of the importance of each explanatory variable in the model. Both importance values were normalized by dividing them by the maximum value for each model. The series of processes was repeated with randomized data splitting only or with both oversampling and data splitting to construct 100 models from the original dataset and 1000 models from the oversampled dataset.

**Figure 2.**
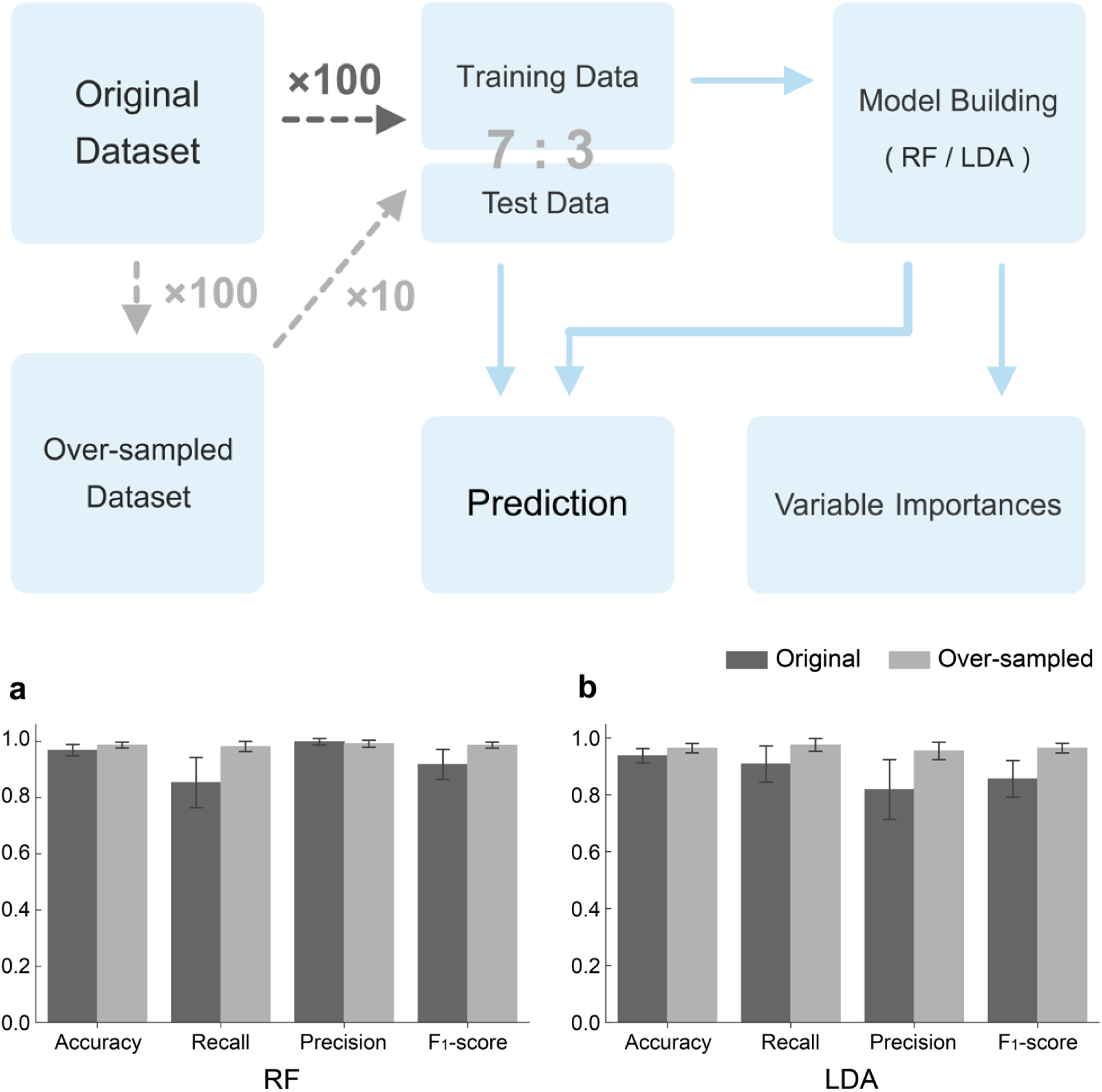
**Procedure of model building (top) and precision (bottom) for the RF (a) and LDA (b) models.**

Details of the datasets and scripts used, as well as the results of all experiments, are available at https://github.com/UTForestChemistryLab/rf-comparative-genomics.

## Results and Discussion

### LDA and RF analysis

LDA calculates the straight line that maximizes the ratio of the within-group variance to the between-group variance of the projection on that line and discriminates the two groups based on which side of the hyperplane each sample lies. The function representing this line (discriminant function) is expressed in the form of a linear combination of the explanatory variables, so that the degree of contribution of each explanatory variable to the discrimination can be read from its coefficient. In contrast, the RF algorithm, proposed by L. Breiman in 2001, follows a concept of ensemble learning called bagging(11) and uses majority voting of a large number of weak learners built in parallel to obtain the final prediction result (Fig. 1). The individual weak learners are simple models, so it is difficult to achieve high accuracy individually, but since they are built from randomly selected data and each is affected by randomness in a different direction, aggregating a large number of weak learners improves the generalization ability and produces a highly accurate model. As a weak learner, we use a decision tree.

RF is a machine learning method with several unique features. First, although data in the life science field generally have a large number of variables and a small number of samples (large p, small n), RF can often achieve high accuracy even in such cases, compared to other algorithms. In addition, the versatility of RF for both classification and regression, the small number of parameters to be adjusted, and the high training speed makes it easy to use even for non-experts. The greatest advantage of RF for use in comparative genomics is that, like LDA, it can calculate the importance of each explanatory variable in the model and throw light on the causal relationship between the explanatory variable (gene) and the target variable (trait). The structure of machine learning models is often a “black box”. However, the focus of comparative genomics is not only to predict traits from genomic data, but also to identify the genes involved in determining those traits. Therefore RF, a machine learning algorithm that can calculate the importance of each explanatory variable and interpret the structure of the data, is an excellent choice for comparative genomics.

### Phylogenetic Analysis and Effects of Oversampling

The decay mode dataset used in the experiment consisted of 232 CAZymes annotated data samples from 15 orders (including two samples of unknown classification) (Table 3). The breakdown of decay modes was 183 samples for white-rot fungi and 49 samples for brown-rot fungi. White-rot fungi were represented by 11 orders, with high percentages of Agaricales and Polyporales fungi (41% and 33% of the total, respectively). Brown-rot fungi, on the other hand, were found in seven orders, with Polyporales accounting for the largest proportion (37%). Most orders were populated by one of the decay styles, and only three orders (Agaricales, Polyporales, and Amylocorticiales) contained both white- and brown-rot fungi.

**Table 3.** Composition of samples included in the decay mode dataset.

| Order | White-rot | Brown-rot |
| --- | --- | --- |
| Agaricales | 75 | 1 |
| Amylocorticiales | 2 | 5 |
| Atheliales | 3 | 0 |
| Auriculariales | 2 | 0 |
| Boletales | 0 | 10 |
| Cantharellales | 1 | 0 |
| Corticiales | 8 | 0 |
| Geastrales | 1 | 0 |
| Gloeophyllales | 0 | 5 |
| Hymenochaetales | 11 | 0 |
| Jaapiales | 0 | 1 |
| Polyporales | 60 | 18 |
| Russulales | 15 | 0 |
| Trechisporales | 3 | 0 |
| Dacrymycetales | 0 | 9 |
| unclassified | 2 | 0 |
| Overall | 183 | 49 |

The phylogenetic tree formed a clade consistent with the NCBI order-level classification, except that the Polyporales and Corticiales were each split into three lineages (Fig. S1). The branch divisions, including the Polyporales and Corticiales clades, were also in good agreement with the phylogenetic tree provided by JGI MycoCosm. Four major groups of brown-rot fungi were identified in the phylogenetic tree, including a clade of Dacrymycetes, but *Leptoporus mollis* (Lepmol1) of Polyporales and *Fistulina hepatica* (Fishe1) of Agaricales were isolated from the other brown-rot fungi and placed alone in the white-rot group.

The LDA and RF models showed high percentages of correct responses (LDA: 94.6%, RF: 96.8%), even when trained on the original imbalanced decay mode dataset. On the other hand, both methods showed divergent values for fit and recall, suggesting a bias in the predictive ability of the models. The models using the oversampled dataset for training showed better predictive accuracy than the models trained on the original dataset for most indicators, for both LDA and RF (Fig. 2). Most importantly, the divergence between goodness of fit and reproducibility was eliminated, and the F1 score, their harmonic mean, was also improved. Indeed, the models trained on the oversampled dataset were able to predict the two decay modes in a balanced manner with high accuracy in both cases. Therefore, both the LDA and RF prediction models were considered to appropriately reflect the differences in CAZymes composition between white- and brown-rot fungi (i.e., properly reflecting the importance of each explanatory variable).

### Cellulose-degrading enzymes contributing to the prediction of decay modes

In the RF model, Gini importance was used as a measure of the importance of each explanatory variable. Gini importance, also called mean impurity reduction, is calculated by averaging across the RF how much the impurity (degree of white/brown-rot fungus admixture) of the child nodes is decreased relative to the parent node in each decision tree in the model when the data were split with respect to that variable.

The most important explanatory variable in the RF model trained on the oversampled dataset in SMOTE was the number of genes in the AA9 family (Fig. 3a). Although the model included 266 CAZy families/subfamilies as explanatory variables, only a small number of them seemed to make a strong contribution to the prediction, with 31 families having an importance greater than 0.1 (one-tenth of one percent for AA9) and 18 families having an importance greater than 0.2 (Table S1). AA9 genes were present in all Agaricomycetes genome samples regardless of the decay mode, but the distribution of AA9 genes diverged by approximately 10 genes between white- and brown-rot fungi. This divergence in distribution may have led to a significant reduction in impurity when nodes were partitioned based on AA9 in the decision tree, increasing the importance of this family (Fig. 3c). AA9 is a family of LPMOs that primarily use cellulose as a substrate. This enzyme oxidatively cleaves β-1,4-glycosidic linkages by hydroxylating the C1 and/or C4 positions of glucose residues on the surface of crystalline cellulose (25–28). The greatest importance of the AA9 family is likely due to the role of oxidative cellulose degradation mechanisms in wood decay. It has also recently been suggested that LPMOs may also be involved in lignin degradation(29), and more detailed elucidation of the function of this family is needed.

**Figure 3.**
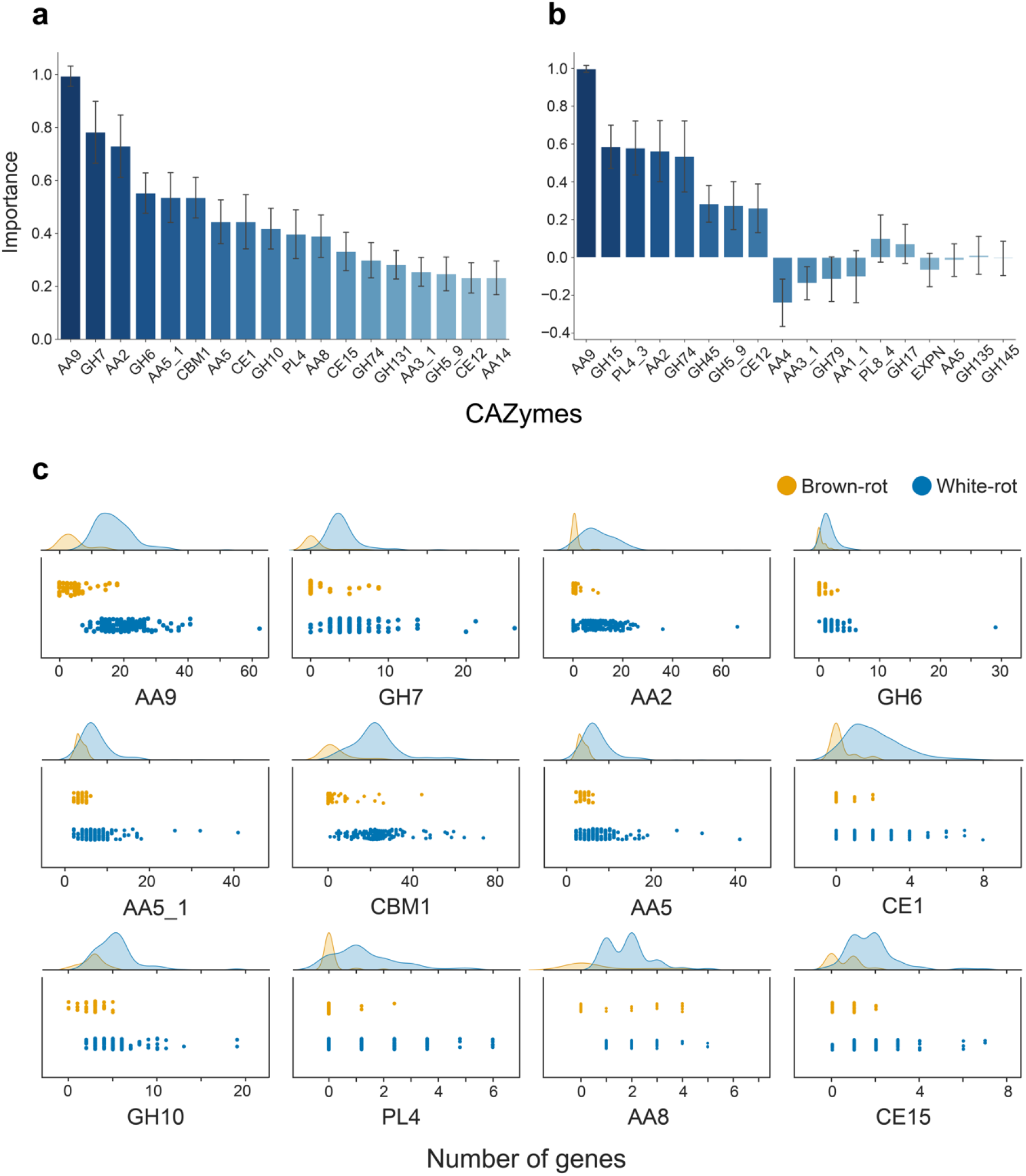
Importance of each CAZy family in the model and distribution of number of genes per decay mode. The importance of each explanatory variable (CAZy family/subfamily) in the RF (a) and LDA (b) models (relative to the family of greatest importance). For RF, only the top 18 families are shown. Error bars represent standard deviations. (c) Distribution of the number of genes in the top 12 most important families in the RF model. A scatter plot (jitter plot) and its kernel density estimation graph are shown.

In addition to AA9, other groups of enzymes that act on crystalline cellulose were of high importance, with GH7 and GH6, the cellulase family including CBH, having the second and fourth highest importance, respectively. CBM1, a cellulose-binding CBM, had the sixth highest importance. More than 80% of brown-rot fungi lacked either GH6 or GH7, and 50% completely lacked any CBH gene. Conversely, most white-rot fungi had at least one gene from each of the two families. Three species of Hymenochaetales, *Phellinus ferrugineofuscus* (Phefer1), *Phellopilus nigrolimitatus* (Pheni1), and *Sidera vulgaris* (Sidvul1), lacked the GH7 gene, and the Polyporales *Ceriporiopsis subvermispora* (Cersu1) lacked the GH6 gene, but no white-rot fungi lacked both GH6 and GH7.

Cellobiose dehydrogenase (CDH), an enzyme that catalyzes the oxidation of cellobiose to cellobionolactone and acts synergistically with LPMO and CBH, was also represented near the top of the importance scale (11th and 15th), as were the two domains AA8 and AA3_1 that make up this enzyme. All these genes were present in at least one or more of the white-rot genomes.

On the other hand, the cellulase family/subfamily consisting only of endoglucanases (EGs) that use amorphous cellulose as a substrate and do not contain CBH, GH45 (0.114: 28th), GH12 (0.040: 55th), GH5_5 (0.021: 79th) or GH 9 (0.013: 113th) were not as important (Table S1). The differences in the number of genes among the decay modes for these cellulose degradation-related families were consistent with previous comparative genomics observations(5, 8, 17).

### Lignin-degrading enzymes

The AA2 family of lignin-degrading enzymes, or class II peroxidases (PODs), was the third most important after AA9 and GH7. AA2 genes were possessed by more than half of the brown-rot fungi, mainly Polyporales, but the number of copies varied greatly, with white-rot fungi possessing on average about 10 times as many AA2 genes as brown-rot fungi.

The AA5_1 subfamily of glyoxal oxidase (GLOX) and the AA5 family, which includes other copper radical oxidases with different substrates, such as galactose and raffinose, also ranked high, at the 5th and 7th places, respectively. All decaying fungi had at least two AA5_1 genes, but the brown-rot fungi had at most six AA5_1 genes, while the white-rot fungi had many more genes, which may have led to a significant reduction of impurity in the node partitioning of the decision tree based on this family. POD requires H_2_O_2_ for its ligninolytic activity, which is thought to be provided by the enzymatic reaction of GLOX(30). The fact that both AA2 and AA5 gene numbers contributed significantly to the prediction of the decay mode suggests that the linkage of these two families plays a role in the ligninolytic system of white-rot fungi. AA3 (glucose-methanol-choline oxidoreductase) and AA7 (glucooligosaccharide oxidase) are also potential sources of H_2_O_2_, but their importance values were different. AA3 and AA7 were about one-fourth (0.123: 26th) and less than one-thirtieth (0.018: 93rd), respectively, as important as AA5. It is possible that the mechanism of H_2_O_2_ supply from these enzyme families to POD is either non-existent or operates only in a limited number of strains.

Surprisingly, the importance of AA1_1, a subfamily of laccase (Lac), which is considered a central lignin-degrading enzyme along with POD, was not so high: its importance was 0.110, less than one-sixth of that of AA2, placing it 30th overall. In fact, there is a large overlap in the distribution of AA1_1 genes between white- and brown-rot fungi (Fig. S3), and many white-rot fungi had no genes in this subfamily at all, while the enzyme was also used by some brown-rot fungi.

### Hemicellulose-degrading enzyme

After cellulose and lignin degrading enzymes, the family of hemicellulose degrading enzymes was also represented near the top in importance. In particular, the family of enzymes involved in xylan degradation was very important, with GH10, a family of xylanases, ranked 9th in importance, followed by CE1 (acetyl xylan esterase: a family of enzymes that help xylanases access their substrates by acting on the xylan side chain). These families, as well as AA5_1, appear to have increased in importance due to the large number of white-rot fungi with more genes than the upper limit of the number of genes in brown-rot fungi. AA14, a family of LPMOs that use xylan as a substrate, also showed a slightly higher importance (0.232: 18th position). On the other hand, the importance of another xylanase family, GH11 (0.083: 33rd position), and the xylan-binding CBM, CBM13 (0.078: 34th position), was not increased much. The number of genes in this family may be related to the host tree species rather than the mode of degradation.

Glucomannan is also the main hemicellulose of secondary cell walls in conifers, together with xylan, but there was no family of mannanases of high importance. Instead, a group of enzymes related to the degradation of primary wall polysaccharides, led by PL4 (rhamnogalacturonan endolyase) in the 10th position, CE12 (rhamnogalacturonan acetyl esterase), a group of enzymes involved in pectin degradation and a family of xyloglucanases, such as GH74 and GH79, showed an importance of more than 0.1. The differences between the white- and brown-rot genomes as regards these primary wall polysaccharides and enzyme families involved in xylan degradation were missed in previous studies, and the ability to detect these subtle differences is considered an advantage of comparative genomics using machine learning.

In addition, all of the CAZy families of highest importance, including these newly implicated degradation mode families, had more genes in white-rot fungus, and no genes were found that characterized brown-rot fungus. This may indicate that the degradation of woody biomass by brown-rot fungi is mediated by a different mechanism that does not rely on CAZymes.

In the LDA model, the major and minor relationships of importance were also broadly consistent. For example, the number of genes in the AA9 family was most important, followed by PL4_3, AA2, and GH74, and the lignin and primary wall polysaccharide degrading enzymes were represented at the top of the list (Fig. 3b). Crystalline cellulose and xylan degrading enzymes were mostly represented by AA9, due to the refinement of explanatory variables during model construction (Table 2). In principle, for the number of genes of a family, if white-rot fungi tend to be more abundant, the importance of that family takes a positive value in LDA, while if they are less abundant, the importance takes a negative value. However, AA3_1 and AA5 had negative importance values, even though the white-rot fungus contained more genes. This is due to a phenomenon called coefficient inconsistency, and arises from the strength of the correlation between the explanatory variables.

Thus, the fact that the values fluctuate depending on the correlations among the explanatory variables, and the need to refine the explanatory variables in advance to minimize their impact, make it difficult to understand the impact of each CAZy family in the LDA model. On the other hand, in the RF model, the independent importance (Gini importance) can be calculated for all the families, independently of the correlations, though the value represents only the degree of contribution to the prediction, not a direct expression of the effect of each explanatory variable on the target variable. In contrast, the importance in LDA is a coefficient of a linear discriminant function, and if one explanatory variable is twice as important as another, this means that the change in the target variable is twice as large when the explanatory variable increases by one. Thus, the fact that the importance in the LDA analysis, which more clearly expresses the causal relationship between explanatory and target variables, matched the trend in the importance of each CAZy family found in the RF analysis is considered to support the reliability of comparative genomics using the RF model.

### Diversity of decay modes

When the confidence values in predicting the test sample to be a white- or brown-rot fungus (discrimination scores for LDA and class probability estimates for RF) calculated by each model over 1000 performance tests were averaged, 11 samples were incorrectly predicted to be white- or brown-rot fungi by LDA, while 7 of these were correctly predicted to be white-rot fungi by RF (Fig. 4).

**Figure 4.**
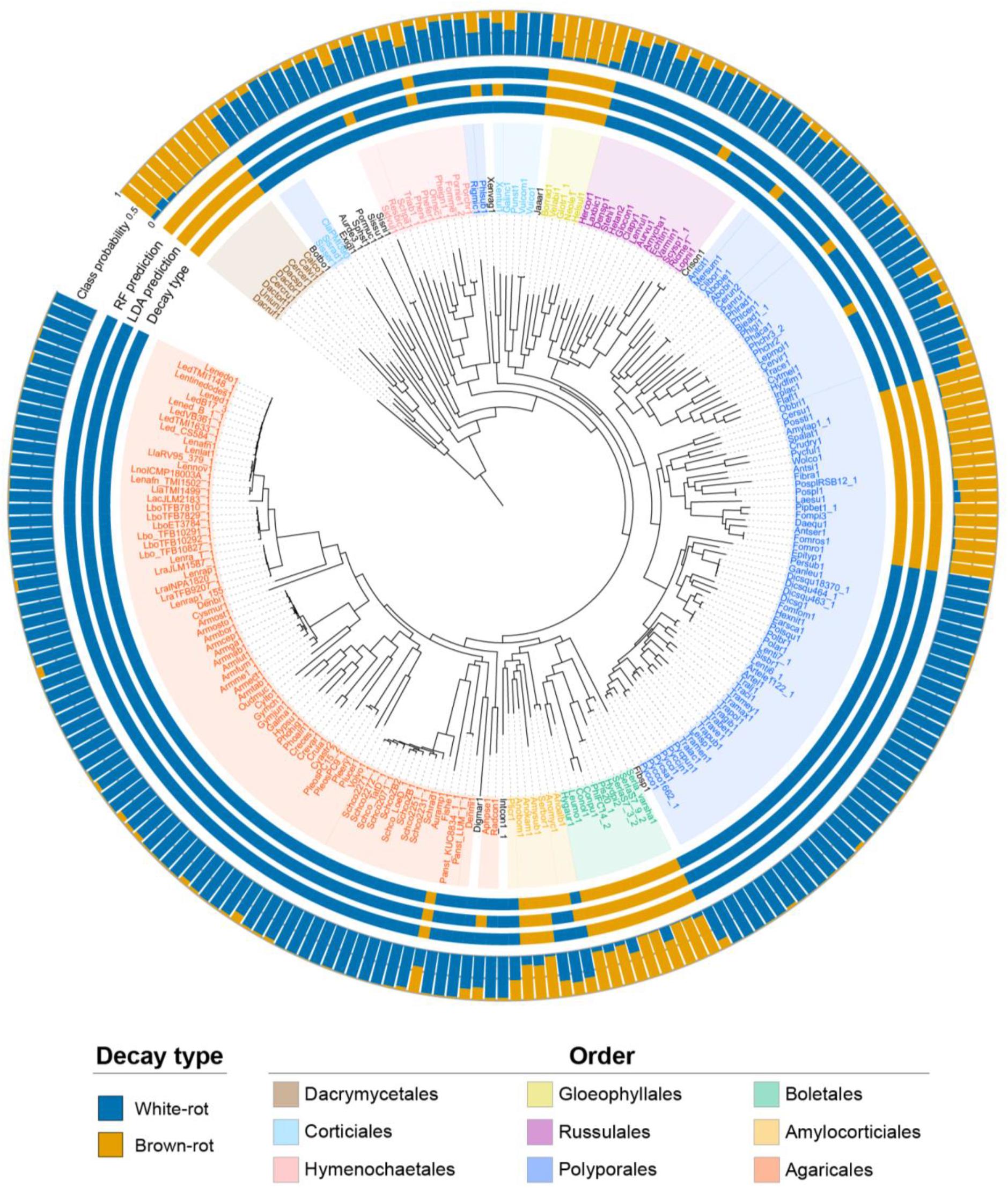
RF prediction results from the model. The correct decay style (Decay type) and the results of prediction by LDA and RF (LDA/RF prediction) for each fungus in the dataset. Prediction results were determined from the average of the level of confidence in the output prediction (LDA: discriminant score, RF: estimate of class probability). The average of the estimates of class probability in RF is also shown in the figure (Class probability).

Of the four samples incorrectly predicted by RF, *Onnia scaura* (Onnsc1) was predicted to be a brown-rot fungus, although it is actually a white-rot fungus and the only closely related species is also a white-rot fungus. This fungus had fewer genes than the average white-rot fungus for the most important CAZymes in the RF model, except for GH10, and its CAZymes composition was closer to the average for brown-rot fungi (Fig. 5).

**Figure 5.**
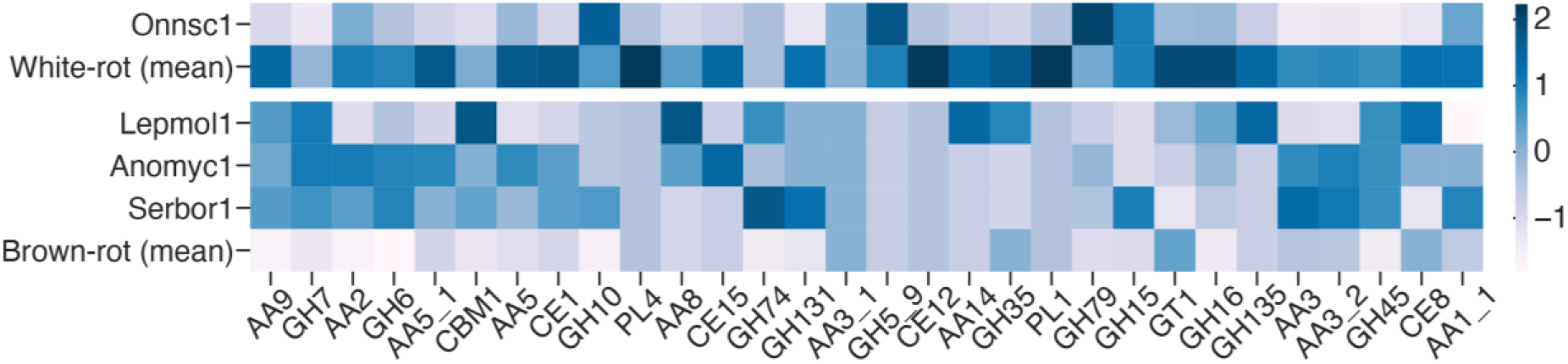
CAZymes Composition of Wrongly Predicted Fungi. Heatmap of the number of CAZymes genes in the four species incorrectly predicted by the RF model compared to the average of white- and brown-rot fungi. Families are the top 30 families in the RF model, and the number of genes is standardized for each family.

However, AA2, AA5 and AA1_1 had more genes than the average brown-rot fungus, although they did not reach the average for white-rot fungus. Conversely, *Leptoporus mollis* (Lepmol1), which is actually a brown-rot fungus but was incorrectly predicted to be a white-rot fungus, had significantly more genes than the brown-rot average (and in some families, more than the white-rot average), mainly among crystalline cellulose-related CAZymes. However, the number of genes in the lignin degradation-related families was not much different from the brown-rot average, especially AA1_1, for which no genes at all were present.

These fungi have CAZymes sets that are typical of white- and brown-rot fungi, respectively, with respect to lignin degradation, but they are mispredicted because of their genome organization, which resembles the opposite degradation mode with respect to other components such as crystalline cellulose. The presence of such fungi suggests the diversity of wood decay systems. On the other hand, the CAZymes compositions of *Anomoporia myceliosa* (Anomyc1) and *Serpulomyces borealis* (Serbor1), which are brown-rot fungi but predicted to be white-rot fungi according to the literature, are related not only to crystalline cellulose degradation, but also to the lignin degradation family, and were close to those of white-rot fungi for both the crystalline cellulose-related and lignin degradation-related families. These two species are both Amylocorticiales, and closely related species are a mixture of white/brown-rot fungi. Other family members, such as xylan degrading enzymes (CE1, GH10, CE15), tended to have more genes than the brown-rot average, although there was some variation between the two species. Thus, these fungi have genomes similar to those of white-rot fungi for almost all CAZymes that may play a central role in determining the decay mode, raising the possibility that previous reports may have been in error. If the results of re-examination confirm that these two species are brown-rot fungi, it is possible that they have a wood decay system that is different from that of other brown-rot fungi.

The RF model correctly predicted all samples except for these four species. For more than 85% of all samples, the predictions were unambiguous, with class probabilities (the probability that the sample belonged to brown-rot fungi) of less than 0.2 or greater than 0.8 (Fig. 6). On the other hand, 34 samples (24 white-rot fungi and 10 brown-rot fungi), including the four incorrectly predicted species, had class probability estimates ranging from 0.2 to 0.8. Among the samples with these less confident predictions were *Botryobasidium botryosum* (Botbo1, class probability = 0.249), previously reported as a gray-rot fungus, and *Jaapia argillacea* (Jaaar1, class probability = 0.632).

**Figure 6.**
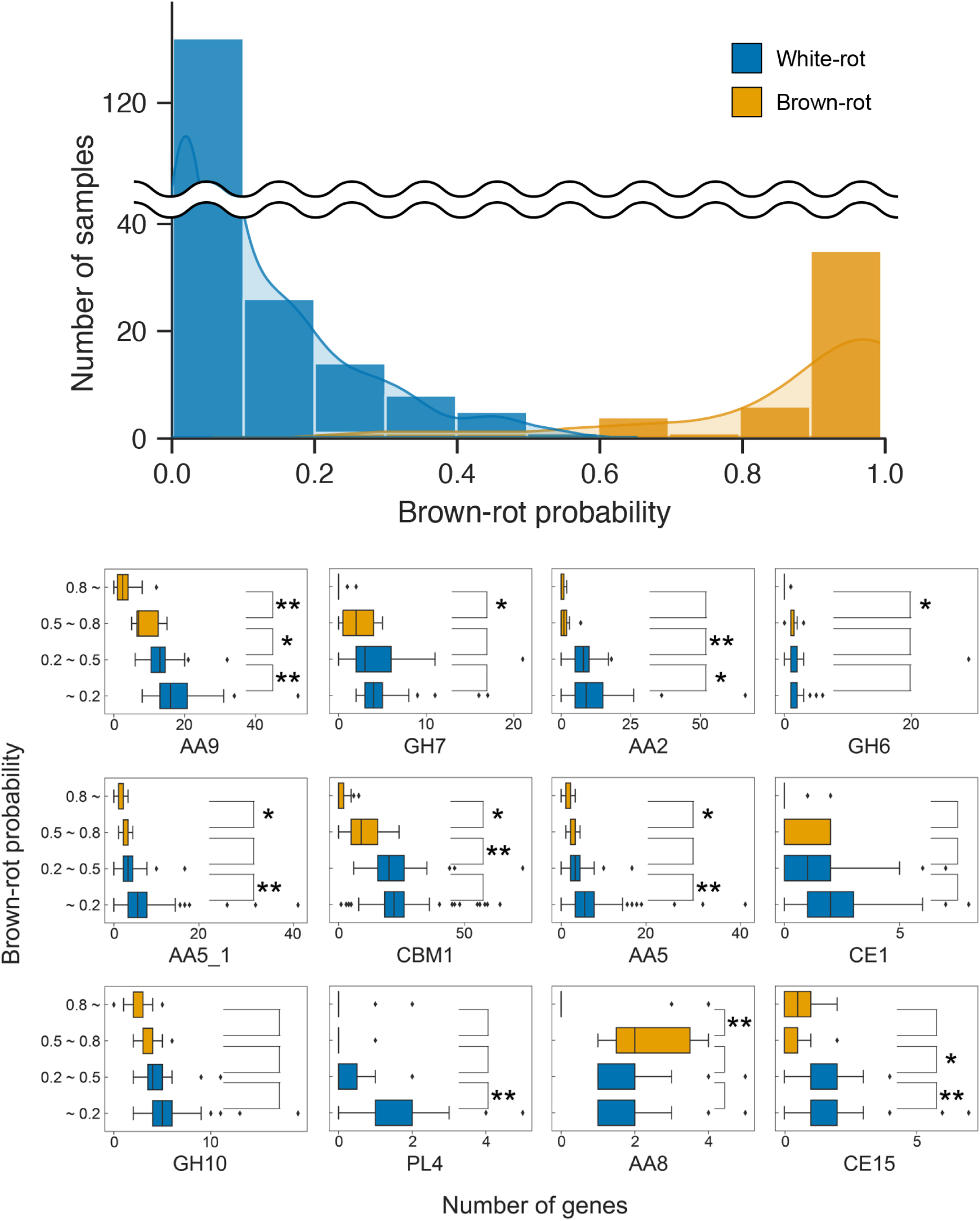
Histogram of estimated class probabilities (top) and distribution of number of genes per class probability estimate (bottom). Each sample in the dataset was grouped according to the predicted results of the RF model (class probability estimates of brown-rot fungi), and the number of genes in the CAZy family with the highest importance for each group is shown in the box-and-whisker plot. Welch’s t-test was performed between adjacent groups at the 5% significance level (*: p < 0.05, **: p < 0.01) under the null hypothesis that “there is no difference in gene numbers between the two groups”.

There was no significant difference in the number of genes between this group of gray-rot fungi and the group of true white-rot fungi (class probability less than 0.2) for crystalline cellulose-related families such as GH7, GH6, AA8, and AA3_1 (Fig. 6, Fig. S4), suggesting that the ability of these fungi to degrade crystalline cellulose is similar to that of true white-rot fungi. However, the number of genes for AA9 was significantly lower, intermediate between those of true white- and brown-rot fungi, suggesting that there may be differences in cellulose degradation mechanisms and abilities depending on the magnitude of the role of LPMO in decay, considering the high importance of this family in predicting the decay mode.

In the lignin degradation-related family, there was a significant difference in the number of genes for AA2 between the gray-rot group, which was closer to the brown-rot group (class probability of 0.5-0.8), and the white-rot group (0.2-0.5). On the other hand, there was no significant difference between the gray-rot group and the true brown-rot group, suggesting that the number of genes for this enzyme may be a critical factor in the ability to degrade lignin. In contrast, there was no significant difference in the number of genes for AA1_1 between the groups, again suggesting that Lac is not a universal factor separating the white-rot/brown-rot decay modes.

The presence of brown-rot fungi with a crystalline cellulolytic enzyme gene set comparable to that of white-rot fungi, and vice versa, in samples that were mispredicted or potentially erroneously assigned as described above, suggests a diversity of decay modes. Analysis of the importance of each explanatory variable also showed that for all CAZymes, including POD and CBH, which are typical genes that separate white- and brown-rot fungi, there was no clear family of boundaries that completely separated the two decay modes based on the presence or number of genes. Therefore, there appears to be no absolute general rule regarding the factors that determine the decay mode of wood-rotting basidiomycetes. The results suggest that wood-rotting fungi use different strategies to decompose woody biomass and that their decay modes cannot be described by a simple dichotomy.

## Conclusion

In the present study, we show that the machine learning RF algorithm can successfully classify white- and brown-rot fungi according to the number of CAZyme families in the basidiomycete genome. The accuracy of the RF algorithm reached 98%, which indicates that the classification method is useful for comparative genomics without human bias. Moreover, the most important enzyme families that divide white- and brown-rot fungi are those involved in crystalline cellulose (AA9 and GH7) and lignin (AA2) degradation, in accordance with previous results of comparative genomics studies. All these families are indispensable for white-rot fungi, which may imply that the degradation mechanism by brown-rot fungus could be either non-enzymatic and/or enzymatic employing an unknown system for wood digestion.

## Acknowledgements

We thank Dr. Akio Nakabayashi at Yokogawa Electric Corporation for technical discussions about the analysis. This study received financial support in the form of Grants-in-Aid for Scientific Research (A) from the Japan Society for the Promotion of Science (JSPS, No. 23H00341) to KI.

## Author contributions

NH contributed to the data analysis and draft writing, MS supported selection of machine learning algorithms, and KI designed the experiments and wrote the manuscript.

## Abbreviations

AA: auxiliary activity; CAZymes: Carbohydrate-Active enZymes; CBH: cellobiohydrolase; CBM: carbohydrate-binding modules; CE: carbohydrate esterase; GH: glycoside hydrolase; GT: glycosyltransferase; LDA: linear discriminant analysis; LPMO: lytic polysaccharide monooxygenase; PL: polysaccharide lyase; POD: class II peroxidase.

## References

1. Horwath, W. (2007) 12 - Carbon cycling and formation of soil organic matter In Soil Microbiology, Ecology and Biochemistry (Third Edition), Paul EA, ed. Academic Press, San Diego 303–339

2. Moon, R. J., Martini, A., Nairn, J., Simonsen, J., and Youngblood, J. (2011) Cellulose nanomaterials review: structure, properties and nanocomposites Chem Soc Rev 40, 3941–3994 10.1039/C0CS00108B

3. Ralph, J., Lapierre, C., and Boerjan, W. (2019) Lignin structure and its engineering Curr Opin Biotechnol 56, 240–249 10.1016/j.copbio.2019.02.019

4. Kirk T K. (1973) Quantitative changes in structural components of conifer woods during decay by white-and brown-rot fungi Phytopathology 63, 1338–1342 10.1094/phyto-63-1338

5. Riley, R., Salamov, A. A., Brown, D. W., Nagy, L. G., Floudas, D., Held, B. W. et al. (2014) Extensive sampling of basidiomycete genomes demonstrates inadequacy of the white-rot/brown-rot paradigm for wood decay fungi Proc Natl Acad Sci USA 111, 9923–9928 10.1073/pnas.1400592111

6. Yelle, D. J., Wei, D., Ralph, J., and Hammel, K. E. (2011) Multidimensional NMR analysis reveals truncated lignin structures in wood decayed by the brown rot basidiomycete Postia placenta Environ Microbiol 13, 1091–1100 10.1111/j.1462-2920.2010.02417.x

7. Lombard, V., Golaconda Ramulu, H., Drula, E., Coutinho, P. M., and Henrissat, B. (2014) The carbohydrate-active enzymes database (CAZy) in 2013 Nucl Acids Res 42, D490-D495 10.1093/nar/gkt1178

8. Floudas, D., Binder, M., Riley, R., Barry, K., Blanchette, R. A., Henrissat, B. et al. (2012) The paleozoic origin of enzymatic lignin decomposition reconstructed from 31 fungal genomes Science 336, 1715–1719 10.1126/science.1221748

9. Kohler, A., Kuo, A., Nagy, L. G., Morin, E., Barry, K. W., Buscot, F. et al. (2015) Convergent losses of decay mechanisms and rapid turnover of symbiosis genes in mycorrhizal mutualists Nat Genet 47, 410–415 10.1038/ng.3223

10. Fisher, R. A. (1936) The use of multiple measurements in taxonomic problems Annal Eugen 7, 179–188 10.1111/j.1469-1809.1936.tb02137.x

11. Breiman, L. (2001) Random forests Machine Learning 45, 5–32 10.1023/A:1010933404324

12. Chen, X., and Ishwaran, H. (2012) Random forests for genomic data analysis Genomics 99, 323–329 10.1016/j.ygeno.2012.04.003

13. Touw, W. G., Bayjanov, J. R., L, O., L, B., J, B., M, W., and Hijum, S. A. F. T. v. (2012) Data mining in the Life Sciences with Random Forest: a walk in the park or lost in the jungle? Brief Bioinformatics 14, 315–326 10.1093/bib/bbs034

14. Grigoriev, I. V., Nikitin, R., Haridas, S., Kuo, A., Ohm, R., Otillar, R., et al. (2014) MycoCosm portal: gearing up for 1000 fungal genomes Nucl Acids Res 42, D699–D704 10.1093/nar/gkt1183

15. Worrall, J. J., Anagnost, S. E., and Zabel, R. A. (1997) Comparison of wood decay among diverse lignicolous fungi Mycologia 89, 199–219 10.1080/00275514.1997.12026772

16. Filippova, N. V., and Zmitrovich, I. V. (2013) Wood decay community of raised bogs in West Siberia Environ Dynam Glob Climate Change 4, 1–16,

17. Nagy, L. G., Riley, R., Tritt, A., Adam, C., Daum, C., Floudas, D. et al. (2016) Comparative genomics of early-diverging mushroom-forming fungi provides insights into the origins of lignocellulose decay capabilities Mol Biol Evol 33, 959–970 10.1093/molbev/msv337

18. Enright, A. J., Van Dongen, S., and Ouzounis, C. A. (2002) An efficient algorithm for large-scale detection of protein families Nucl Acids Res 30, 1575–1584 10.1093/nar/30.7.1575

19. Katoh, K., Rozewicki, J., and Yamada, K. D. (2019) MAFFT online service: multiple sequence alignment, interactive sequence choice and visualization Brief Bioinformatics 20, 1160–1166 10.1093/bib/bbx108

20. Gouy, M., Guindon, S., and Gascuel, O. (2010) SeaView Version 4: a multiplatform graphical user interface for sequence alignment and phylogenetic tree building Mol Biol Evol 27, 221–224 10.1093/molbev/msp259

21. Capella-Gutiérrez, S., Silla-Martínez, J. M., and Gabaldón, T. (2009) trimAl: a tool for automated alignment trimming in large-scale phylogenetic analyses Bioinformatics 25, 1972–1973 10.1093/bioinformatics/btp348

22. Sánchez, R., Serra, F., Tárraga, J., Medina, I., Carbonell, J., Pulido, L. et al. (2011) Phylemon 2.0: a suite of web-tools for molecular evolution, phylogenetics, phylogenomics and hypotheses testing Nucl Acids Res **39**, W470–W474 10.1093/nar/gkr408

23. Trifinopoulos, J., Nguyen, L.-T., von Haeseler, A., and Minh, B. Q. (2016) W-IQ-TREE: a fast online phylogenetic tool for maximum likelihood analysis Nucl Acids Res 44, W232–W235 10.1093/nar/gkw256

24. Chawla, N. V., Bowyer, K. W., Hall, L. O., and Kegelmeyer, W. P. (2002) SMOTE: synthetic minority over-sampling technique J Artif Int Res 16, 321–357,

25. Beeson, W. T., Phillips, C. M., Cate, J. H. D., and Marletta, M. A. (2012) Oxidative cleavage of cellulose by fungal copper-dependent polysaccharide monooxygenases J Amer Chem Soc 134, 890–892 10.1021/ja210657t

26. Horn, S. J., Vaaje-Kolstad, G., Westereng, B., and Eijsink, V. (2012) Novel enzymes for the degradation of cellulose Biotechnol Biofuels 5, 45 10.1186/1754-6834-5-45

27. Isaksen, T., Westereng, B., Aachmann, F. L., Agger, J. W., Kracher, D., Kittl, R. et al. (2014) A C4-oxidizing lytic polysaccharide monooxygenase cleaving both cellulose and cello-oligosaccharides J Biol Chem 289, 2632–2642 10.1074/jbc.M113.530196

28. Uchiyama, T., Uchihashi, T., Ishida, T., Nakamura, A., Vermaas, J. V., Crowley, M. F., et al. (2022) Lytic polysaccharide monooxygenase increases cellobiohydrolases activity by promoting decrystallization of cellulose surface Science Adv 8, eade5155 10.1126/sciadv.ade5155

29. Li, F., Ma, F., Zhao, H., Zhang, S., Wang, L., Zhang, X., and Yu, H. (2019) A lytic polysaccharide monooxygenase from a white-rot fungus drives the degradation of lignin by a versatile peroxidase Appl Environ Microbiol **85**, e02803–02818 10.1128/AEM.02803-18

30. Dashtban, M., Schraft, H., Syed, T. A., and Qin, W. (2010) Fungal biodegradation and enzymatic modification of lignin Int J Biochem Mol Biol 11, 36–50,

